# *Pfcrt* copy number amplification detected in a *Plasmodium falciparum* outbreak

**DOI:** 10.64898/2026.06.18.732750

**Authors:** Anjana Rai, Chiyun Lee, Hidayat Trimarsanto, Ashley Osborne, Kian Soon Hoon, Jacob AF Westaway, Giri S Rajahram, June Haidee Acuña-Lariosa, Maria Lourdes M Macalinao, Jennifer Luchavez, Dave Tangcalagan, Sherwin Galit, Anisah Jantim, Kim A Piera, Fe Esperanza Espino, Timothy William, David Fidock, Dominic P Kwiatkowski, Nicholas M Anstey, Bridget E Barber, Richard D Pearson, Matthew J Grigg, Sarah Auburn

## Abstract

Antimalarial resistance is one of the greatest threats to malaria elimination. Following an outbreak of *Plasmodium falciparum* infection in Sabah Malaysia in 2019, 98 samples were sequenced to search for adaptations driving the outbreak. The genomic data revealed evidence of clonal expansion of a strain carrying multiple copies of the chloroquine resistance transporter gene (*pfcrt*) in 86 cases. A TaqMan qPCR assay was developed to quantify *pfcrt* copy number, confirming the *in silico* evidence of duplication. Whilst point mutations in *pfcrt* have been associated with resistance to chloroquine and other drugs, copy number amplification has not been widely explored as a resistance mechanism. We investigated the genetic architecture of the duplication, revealing a wildtype haplotype (3D7 reference-type with 76K variant) and a novel mutant (with 76T mutation). Application of the TaqMan assay in 43 *P. falciparum* cases from neighbouring Palawan Island, the Philippines, identified a further two cases with *pfcrt* duplication. Assessment of the global genomic data in the MalariaGEN Pf8 repository identified a further 86 cases, including 73 (85%) from West Africa with evidence of *pfcrt* duplication. Amongst 47 monoclonal MalariaGEN cases, majority (95%) comprised wild type and mutant variants at codon 76. Our study reveals a potential previously unconsidered antimalarial resistance mechanism for *P. falciparum* and provides an assay for surveillance in other populations.

## Introduction

*Plasmodium falciparum* is the major global cause of malaria morbidity and mortality (WHO, 2025b). Despite initial progress in reducing incidence following calls for the eradication of malaria, case numbers have stalled and even increased in some endemic regions since 2016 (WHO, 2025b). Chemotherapeutic agents have historically played a pivotal role in managing malaria, but the emergence of drug-resistant lineages has undermined their effectiveness^2,1^. This resistance is driven by both the high parasite burden and the inherent mutability of the parasite’s genome^3^. Information on parasite drug resistance mechanisms is critical for surveillance and associated treatment response, as well as for drug development. However, the molecular determinants and associated mechanisms of *P. falciparum* resistance to several widely used antimalarials, such as amodiaquine (AQ) and lumefantrine (LF), are not fully understood (*WHO compendium*) (Niare et al., 2025; WHO, 2025a).

The long half-lives of AQ and LF, 3 to 6 days and 9 to 18 days, respectively, make them highly effective partner drugs in artemisinin-based combination therapies (ACTs), ensuring the clearance of any remaining parasites following cessation of artemisinin (Peto et al., 2022). However, this process can effectively leave AQ and LF to work as monotherapies in the presence of slow clearing artemisinin-resistant parasite lineages or when used in high transmission regions where the risk of exposure and reinfection is elevated. Polymorphisms in the *P. falciparum* chloroquine resistance transporter (*pfcrt*) and *P. falciparum* multidrug resistance 1 (*pfmdr1*) genes, as well as amplification of *pfmdr1*, have been linked with the treatment failure of AQ and LF (Okell et al., 2018; Venkatesan et al., 2014). Despite this link, the exact mechanism of action for resistance has not been confirmed for either drug. Unlike with AQ, LF has never officially been supplied as a monotherapy, only in conjunction with artemisinin, inadvertently inhibiting our ability to conclusively identify LF resistant field isolates (Hastings & Ward, 2005). In resistant parasites, mutations in *pfcrt* are believed to enable the efflux of drug molecules out of the digestive vacuole (DV), away from the haem target, while mutations in the membrane transporter of *pfmdr1* may enable the transport of drugs, including LF, into the DV and away from their primary action site (Beshir et al., 2010; Folarin et al., 2011; Wicht et al., 2020). The *pfmdr1* N86Y variant, specifically, has been identified to be selected for by use of AQ, however the same variant is selected against by therapy with artemether-lumefantrine (AL). Wild-type (WT) parasites were found to be associated with lower *ex vivo* lumefantrine susceptibility compared to N86Y mutant parasites (Yeka et al., 2016).

Despite reporting no indigenous malaria cases due to human-only *Plasmodium* species infections since 2018, Malaysia remains susceptible to imported transmission. In 2019, an outbreak of *P. falciparum* infections occurred in the setting of Banggi Island, in the northeast point of Sabah, East Malaysia. From 2009, artemisinin-based combination therapy (Artemether-lumefantrine [AL] or artesunate-mefloquine [AS-MQ], predominantly the former) replaced chloroquine (CQ) plus sulfadoxine-pyrimethamine (SP) as first-line treatment for uncomplicated *P. falciparum* infections in Malaysia. However, CQ was still available in the community for treatment of other species, such as *P. vivax* and zoonotic *P. knowlesi* Infections. We conducted Illumina whole genome sequencing of *P. falciparum* patient samples collected within the Banggi outbreak to identify potential parasite genetic drivers. Our genomic investigation revealed evidence of a large clonal expansion of a strain comprising multiple copies of the *pfcrt* gene. Further investigation in global parasite populations detected *pfcrt* duplication in a range of countries, with notably high occurrence in west Africa.

## Results

### High quality genetic and genomic data from Malaysia

A flow chart outlining the number of samples, their geographic attribution and data source, for each of the genetic measures applied in the study is presented in Supplementary Figure 1, with individual listings in Supplementary Data 1. Amongst 109 PCR-positive *P. falciparum* cases from the 2019 Banggi Island outbreak that were subjected to whole genome sequencing (WGS), 98 (90%) were successfully sequenced, resulting in 80 (73%) high-quality WGS samples. A total of 97 (89%) of the Banggi outbreak cases had high-quality amplicon sequencing data at the SpotMalaria 100-SNP neutral barcode, with an overlap of 80 cases that had both high-quality WGS and amplicon data. For temporal analyses, additional Malaysian data was available from 333 cases with high-quality WGS data (320 with high-quality SpotMalaria barcode data) collected from state-wide surveys in Sabah Malaysia between 2010-2017. A selection of 300 additional samples collected between 1996-2022 with high-quality WGS data and deriving from 26 other malaria-endemic countries representing the major global regions was retrieved from the MalariaGEN Pf8 open-access repository to support geospatial mapping. Although WGS data was only available for one case from the Philippines, 39 samples from Palawan Island collected in 2013-18 had high-quality amplicon sequencing data at the SpotMalaria barcode and were also included for geospatial mapping.

### Clonal expansion dynamic in the Malaysian P. falciparum outbreak

Using the *F*_WS_ measure on the WGS data, the majority (79%, 63/80) of the Banggi Island outbreak cases were monoclonal, indicative of limited superinfection (Supplementary Table 1, Supplementary Figure 2). Genetic distance analysis revealed that 100% (63/63) of the monoclonal cases had near-identical haplotypes (genetic distance < 0.01), suggestive of the rapid expansion of a single strain (Figure 1a). Neighbour-joining analyses across the full set of 333 Malaysian WGS samples (2010-2019) revealed the 2019 cluster as a distinct event, with additional clonal expansions in the *P. falciparum* population in the decade preceding its elimination in 2018 (Figure 1a). A total of 4 clonal expansions comprising 10 or more cases were detected, with some strains persisting >3 years, indicative of limited outcrossing in a bottlenecking population.

**Figure 1.**
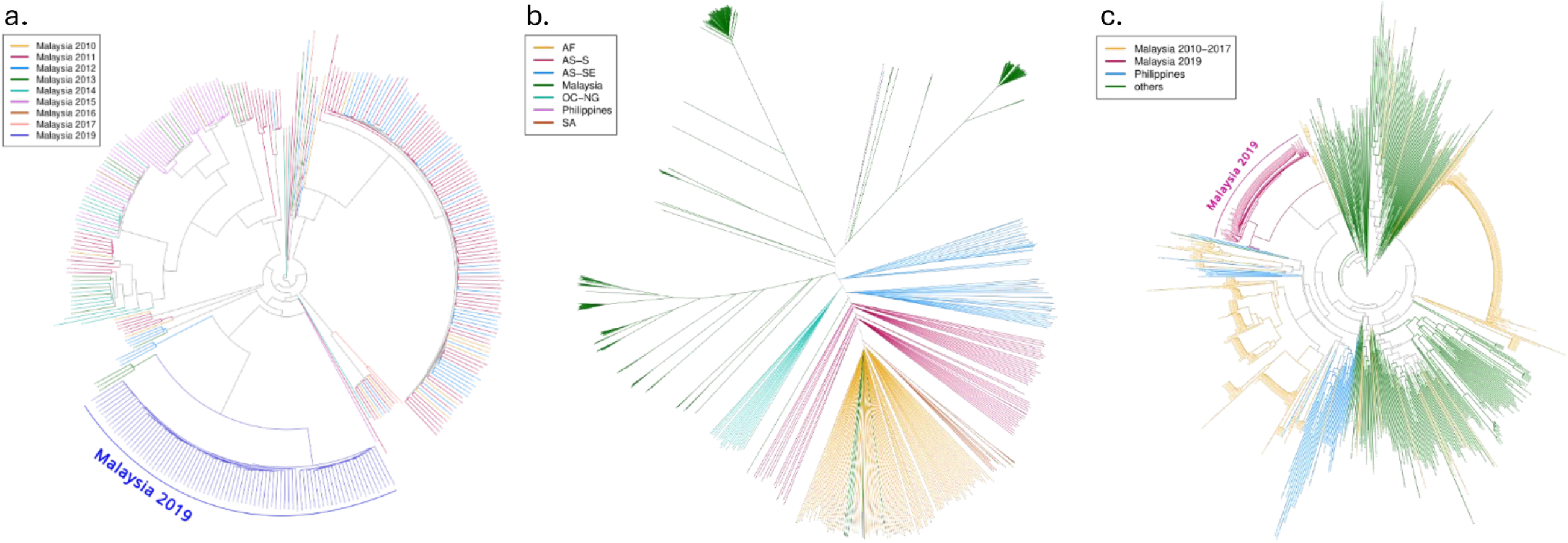
Clonal expansion of an outbreak strain in Malaysian Borneo in 2019. Leaves of the Neighbor-Joining (NJ) trees are colour-coded by sample origin and/or year of collection. Branches are coloured to match their descendant leaves or internal nodes. Where branches of different colours converge, the resulting branch is shown in grey. Panel a) presents a NJ tree of Malaysian samples (N=333) using 116,704 bi-allelic SNPs from WGS data with temporal color representation. The imported/post-elimination Malaysian 2019 samples from Pulau Banggi (N=80) cluster into a single clonal expansion node (dark violet) without any other Malaysian samples in the cluster as indicated by the non-existence of any grey internal branch in the cluster. Panel b) presents a NJ tree of Malaysian (2010-2019) samples together with select samples from the MalariaGEN open-access global data (AF: Africa, AS: Asia, OC-NG: Oceania, SA: South America) including a single Filipino sample. The tree was generated using WGS data at 116,704 bi-allelic SNPs using major alleles at heterozygote positions. All Malaysian samples (dark green) are separate from non-Malaysian samples, regardless of year of sampling, with distinct clusters showing clonal expansions characteristic of a malaria elimination setting. The single Filipino sample (magenta) clusters with the non-clonal Malaysian samples. Panel c) presents a radial nj tree of Malaysian and Filipino samples with temporal-spatial color representation. The tree was generated using the SpotMalaria barcode variants as the samples from the Philippines only had amplicon sequencing data at the barcode. The tree was generated by subsetting WGS data to the SpotMalaria variants and collating this with SpotMalaria amplicon sequencing data. For Malaysian samples with both WGS and amplicon data, the barcode SNPs from WGS data were used. The imported/post-elimination Malaysian 2019 outbreak strain is not observed amongst the Philippines cases but is closely related to them.

Further analysis was conducted including the 300 worldwide cases to determine whether the outbreak strain was local to Sabah or imported from a more distal location. Neighbour-joining analyses using the global data set (n= 732) revealed that the outbreak strain had greatest clustering with other strains from Sabah, suggestive of a local origin (Figure 1b).

Owing to frequent people movement between Sabah and neighbouring Balabac Island (in the province of Palawan) in the Philippines, the potential for a Palawan origin of the Banggi outbreak was investigated using the 39 Palawan cases with SpotMalaria barcode data. Genotype calls at the barcode were successfully derived from amplicon sequencing or WGS data for 431 Sabah and 270 other (global) samples. Neighbour-joining analyses using the barcode data revealed that the outbreak strain had relatively high identity-by-state (IBS) with Palawan compared to other countries (Figure 1c).

### Detection of pfcrt copy number amplification extending >100kb in the post-elimination Sabah outbreak cases

Using the amplicon sequencing and WGS data from the 2019 Sabah cases, we examined WHO-defined validated and candidate drug resistance targets to identify any putative drug resistance or diagnostic evasion variants that could be drivers of the Banggi outbreak. A full list of the variants observed in Sabah in 2019 and other years assessed is presented in Supplementary Data 2. Aside from *pfcrt* and several candidate markers of pyrimethamine and sulfadoxine-pyrimethamine resistance, all of the variants observed in 2019 were wild-type (Table 1). Within *pfcrt*, an excess (>88%) of heterozygote calls were observed at codons 72, 76, 160, 166, 273 and 326 in the outbreak cases (Table 1, Supplementary Figure 3A). This heterozygote pattern was unexpected given the high proportion of monoclonal cases in the outbreak. Computing within-sample allele frequency (WSAF) across all chromosomes in each sample revealed a dense enrichment of heterozygous SNPs in the region surrounding *pfcrt* from Pf3D7_07_v3:403,612-507,272. This region starts within *pfcrt* and ends in the internal hypervariable region of chromosome 7 (Pf3D7_07_v3:508,361-605,650) spanning 25 genes (Supplementary Figure 3B). The presence of region-specific heterozygote calls in evidently monoclonal infections was postulated to reflect copy number amplification encompassing *pfcrt*. Manual inspection of data generated by applying the MalariaGEN Pf8 copy number variant (CNV) pipeline revealed elevated copy number for the same region in some samples, though the signal was insufficient to conclusively characterise the amplification across all suspected samples. In samples where the outputs from the coverage-based analysis were of higher quality, the data suggested the breakpoint was between 401,000 and 403,000, possibly in the intergenic region between *pfsco1* and *pfcrt* (Supplementary Figure 3C). Breakpoints were searched manually using IGV, through custom algorithms to search for potential trans-chromosomal read pairs or read pairs with inappropriate insert sizes, and through BLASTing suspicious softclips of reads, but breakpoints were not found.

**Table 1.** Summary of *validated and candidate drug resistance markers* in the Malaysian outbreak cases. *Prop. Alt. Allele = Proportion of samples carrying an alternate allele with a read depth ≥5; No. Het. = Number of samples with heterozygous alleles at that position; Genotyping source = variant detected using whole genome sequencing (WGS) data and/or SpotMalaria derived amplicon sequencing data. Number of samples (N Samples) with a minimum read depth ≥5 at variant position; Number of samples with a heterozygous call at the variant position (N Het.). Antimalarials include chloroquine (CQ), amodiaquine (AQ), artemisinin (ART), pyrimethamine (PYR), sulfadoxine-pyrimethamine (SP).*Haplotypes could not be accurately derived owing to heterozygote calls.

| Gene | Variant | Prop. Alt. Allele | No. Samples | No. Het. | Drug Assoc. | WHO Status | Genotyping source |
| --- | --- | --- | --- | --- | --- | --- | --- |
| ***Pfcr*** * | C72S | 97% | 189 | 168 | CQ | Validated | WGS, SpotMalaria |
|  | K76T | 97% | 189 | 168 | CQ | Validated | WGS, SpotMalaria |
|  | I218F | 0.5% | 190 | 1 | PPQ | Validated | WGS, SpotMalaria |
|  | H273N | 91% | 189 | 81 | - | - | WGS, SpotMalaria |
|  | N326D | 98% | 191 | 171 | - | - | WGS, SpotMalaria |
| ***Pfmdr***1 | N86Y | 0% | 183 | 0 | CQ, AQ | Validated, Candidate | WGS, SpotMalaria |
| ***Pfk***13 | G687E | 1% | 183 | 2 | - | - | WGS, SpotMalaria |
|  | A675V | 0% | 183 | 0 | ART | Validated | WGS, SpotMalaria |
|  | R622I | 0% | 183 | 0 | ART | Validated | WGS, SpotMalaria |
|  | C580Y | 0% | 183 | 0 | ART | Validated | WGS, SpotMalaria |
|  | P574L | 0% | 183 | 0 | ART | Validated | WGS, SpotMalaria |
|  | V568G | 0% | 183 | 0 | ART | Candidate | WGS, SpotMalaria |
|  | R561H | 0% | 183 | 0 | ART | Validated | WGS, SpotMalaria |
|  | P553L | 0% | 183 | 0 | ART | Validated | WGS, SpotMalaria |
|  | I543T | 0% | 183 | 0 | ART | Validated | WGS, SpotMalaria |
|  | R539T | 0% | 183 | 0 | ART | Validated | WGS, SpotMalaria |
|  | N537I | 0% | 183 | 0 | ART | Candidate | WGS, SpotMalaria |
|  | G538V | 0% | 183 | 0 | ART | Candidate | WGS, SpotMalaria |
|  | G533S | 0% | 183 | 0 | ART | Validated | WGS, SpotMalaria |
|  | R515G | 1.6% | 184 | 3 | - | - | WGS, SpotMalaria |
|  | Y493H | 0% | 184 | 0 | ART | Validated | WGS, SpotMalaria |
|  | M476I | 0% | 184 | 0 | ART | Validated | WGS, SpotMalaria |
|  | C469Y | 0% | 184 | 0 | ART | Validated | WGS, SpotMalaria |
|  | N458Y | 0% | 184 | 0 | ART | Validated | WGS, SpotMalaria |
|  | F446I | 0% | 184 | 0 | ART | Validated | WGS, SpotMalaria |
|  | P441L | 0% | 184 | 0 | ART | Candidate | WGS, SpotMalaria |
|  | E294K | 1% | 88 | 1 | - | - | WGS |
|  | E252Q | 0% | 87 | 0 | ART | Candidate | WGS |
| ***Pfdhfr*** | C59R | 100% | 184 | 0 | - | - | WGS, SpotMalaria |
|  | S108N | 100% | 91 | 0 | - | - | WGS |
|  | S108T | 51% | 96 | 26 | - | - | SpotMalaria |
|  | N51I + S108N | 0% | 88 | 0 | PYR | Candidate | WGS |
|  | C59R + S108N | 100% | 88 | 0 | PYR | Candidate | WGS |
|  | N51I + C59R + S108N | 0% | 88 | 0 | PYR | Candidate | WGS |
| ***Pfdhfr + Pfdhps*** | A581G | 100% | 183 | 0 | - | - | WGS, SpotMalaria |
|  | C59R + S108N + A581G | 100% | 88 | 0 | SP | Candidate | WGS |
|  | C59R + S108N + A437G | 100% | 88 | 0 | SP | Candidate | WGS |
|  | C59R + S108N + A437G + A581G | 100% | 88 | 0 | SP | Candidate | WGS |

Manual IGV inspection was also done to assess the 2019 Sabah samples for amplifications of *pfmdr1* (Pf3D7_05_v3: 955,955 - 963,095), implicated in reduced susceptibility to mefloquine and potentially lumefantrine, as well as *pfpm2* and *pfpm3* (Pf3D7_14_v3 :292,244 - 295,261; Pf3D7_14_v3: 296,683 - 299,101), associated with resistance to piperaquine (WHO Compendium Reference?). No evidence of *pfmdr1*, *pfpm2,* or *pfpm3* copy number amplification was observed, but only 10, 87, and 13 cases, respectively, had sufficient read coverage for investigation within the respective genes and surrounding regions.

To explore whether cases presenting in Sabah in earlier years or cases from neighbouring Palawan Island, the Philippines, had evidence of *pfcrt* amplification, we investigated all available SpotMalaria barcode data from these cases for evidence of excess heterozygotes in *pfcrt.* Fourteen monoclonal cases from Sabah collected between 2010 and 2017 (five from 2012, four from 2013, one from 2015) and one importation from the Philippines in 2011 had a *pfcrt* heterozygote signature at codons 72 and 76. Evaluation of 39 monoclonal samples from Palawan collected between 2013 and 2018 identified a further two cases with a similar *pfcrt* heterozygote signature at codons 72 and 76 suggestive of *pfcrt* amplification (Supplementary Table 2).

A gDNA-based TaqMan real-time PCR assay targeting *pfcrt* was established to determine copy number at the locus. The assay was conducted in 67 cases, including 20 monoclonal cases with the heterozygote *pfcrt* signal: 17 from Sabah (2 from 2013, 15 from 2019) and 3 from the Philippines (1 from 2011, 2 from 2013-2018). A greater than 2-fold (range 1.7 - 2.3-fold) ratio of *pfcrt* to *b-tubulin* gDNA quantity was observed in 20 (100%) of the cases with the heterozygote signature, supporting *pfcrt* copy number amplification (Figure 2, Supplementary Data 3).

**Figure 2.**
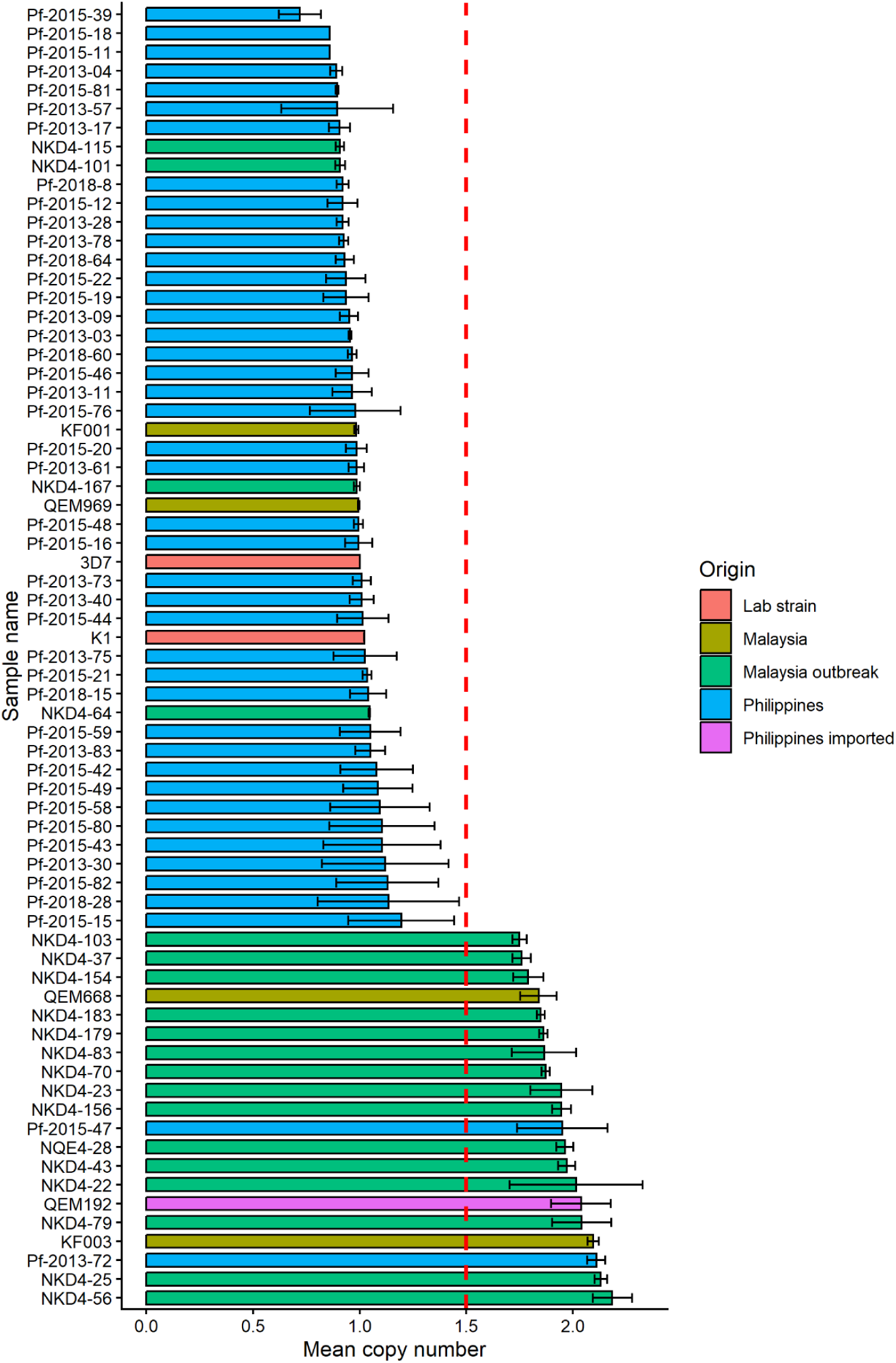
*Pfcrt* copy number estimation in Malaysia and the Philippines by qPCR. Relative copy number of *pfcrt* was assessed via TaqMan qPCR assay using gDNA extracted from malaria patient blood samples. Data are shown as mean (±SD) relative expression of n≥5 replicates. Graph generated using ggplot2, each colour represents the origin of samples.

### A combination of wild type and mutant pfcrt haplotypes in the amplification

Whilst the K76T mutation is well known as the principal determinant of resistance to chloroquine and related aminoquinolines, other *pfcrt* mutations have demonstrated drug modulatory impacts (Petersen et al., 2015). Further exploration of the haplotypic background of the *pfcrt* copies was undertaken using long-read single molecule sequencing (Oxford Nanopore Technologies). Nanopore sequencing of 60 cases, including 19 heterozygotes (16 from Sabah and 3 from the Philippines), revealed two haplotypes, one wild-type haplotype and one novel mutant haplotype comprising codon variants C72S, K76T, A144T, L160Y, I166V and N326D in approximate 1:1 ratios in all 16 heterozygote cases from Sabah and 1 from the Philippines (Table 2, Supplementary Table 3). Two cases from the Philippines had different haplotype combinations from the Sabah duplicates: one comprised two distinct haplotypes both carrying wild-type variants at codon 76, and the second (the importation to Sabah from 2011) comprised one wild-type haplotype and one novel haplotype comprising variants A220S, T333A and I356L (Table 2). Haplotype details for all 79 cases, combined with real-time PCR assay results, are presented in Supplementary Table 3.

**Table 2.** Summary of *Pfcrt* haplotypes observed in Malaysia and the Philippines. The variants within the given haplotypes were derived from Oxford Nanopore Technology (ONT) sequencing data. Shaded variants (besides 166) reflect validated, candidate, or potential determinants of resistance to chloroquine or other antimalarials. *Including the post-elimination 2019 cases. ** single-copy *pfcrt* with minor sequence variation; 1 sample is probably polyclonal with asymmetric haplotype ratio ***Imported case to Malaysia (2011), with travel history to the Philippines.

| Country | No. gene copies | 72 | 73 | 74 | 75 | 76 | 93 | 97 | 101 | 141 | 144 | 145 | 148 | 160 | 166 | 194 | 196 | 218 | 220 | 256 | 271 | 273 | 295 | 326 | 333 | 343 | 353 | 356 | 366 | 367 | 370 | 371 | Haplotype | No. Samples |
| --- | --- | --- | --- | --- | --- | --- | --- | --- | --- | --- | --- | --- | --- | --- | --- | --- | --- | --- | --- | --- | --- | --- | --- | --- | --- | --- | --- | --- | --- | --- | --- | --- | --- | --- |
| *Malaysia | 1 | C | V | M | N | K | T | H | C | V | A | F | L | L | I | I | L | I | A | T | Q | H | N | N | T | M | G | I | A | G | V | R | H1a | 1 |
|  | 1 | S | V | M | N | T | T | H | C | V | T | F | L | Y | V | I | L | I | A | T | Q | H | N | D | T | M | G | I | A | G | V | R | H2 | 7 |
|  | 2 | C | V | M | N | K | T | H | C | V | A | F | L | L | I | I | L | I | A | T | Q | N | N | N | T | M | G | I | A | G | V | R | H1b | 15 |
|  |  | S | V | M | N | T | T | H | C | V | T | F | L | Y | V | I | L | I | A | T | Q | H | N | D | T | M | G | I | A | G | V | R | H2 |  |
| Philippines | 1 | C | V | M | N | K | T | H | C | V | A | F | L | L | I | I | L | I | A | T | Q | H | N | N | T | M | G | I | A | G | V | R | H1a | 25 |
|  | 1 | C | V | M | N | K | T | H | C | V | A | F | L | L | I | I | L | I | A | T | Q | N | N | N | T | M | G | I | A | G | V | R | H1b | 7 |
|  | 1 | C | V | M | N | K | T | H | C | V | A | F | L | L | I | I | L | I | A | T | Q | H | N | N | T | M | G | I | A | G | V | R | H1a | 2** |
|  |  | C | V | M | N | K | T | H | C | V | A | F | L | L | I | I | L | I | A | T | Q | N | N | N | T | M | G | I | A | G | V | R | H1b |  |
|  | 2 | C | V | M | N | K | T | H | C | V | A | F | L | L | I | I | L | I | A | T | Q | N | N | N | T | M | G | I | A | G | V | R | H1b | 1 |
|  |  | C | V | M | N | K | T | H | C | V | T | F | L | L | V | I | L | I | A | T | Q | H | N | D | T | M | G | I | A | G | V | R | H3 |  |
|  | 2 | C | V | M | N | K | T | H | C | V | A | F | L | L | I | I | L | I | A | T | Q | H | N | N | T | M | G | I | A | G | V | R | H1a | 1 |
|  |  | S | V | M | N | T | T | H | C | V | T | F | L | Y | V | I | L | I | A | T | Q | H | N | D | T | M | G | I | A | G | V | R | H2 |  |
|  | 2 | C | V | M | N | K | T | H | C | V | A | F | L | L | I | I | L | I | A | T | Q | N | N | N | T | M | G | I | A | G | V | R | H1b | 1*** |
|  |  | S | V | M | N | T | T | H | C | V | A | F | L | L | I | I | L | I | S | T | Q | H | N | D | A | M | G | L | A | G | V | R | H4 |  |

### High frequency of pfcrt copy number amplification in west Africa with evidence of multiple breakpoints

Using WGS data from the MalariaGEN Pf8 repository, we sought evidence of *pfcrt* copy number amplification in *P. falciparum* cases from geographic regions outside of Sabah and the Philippines. Among 86 samples in Pf8 with evidence of *pfcrt* amplification, the majority came from west Africa, with amplifications observed in seven different countries. The largest number of cases with *pfcrt* amplification was observed in Ghana and these contributed 0.89% of all cases tested in the country (39/4366) (Figure 3a, Table 3). The highest frequency of *pfcrt* amplified cases was observed in Burkina Faso (9/53, 17%) followed by Mali (20/1148, 1.74%) and Côte D’Ivoire (1/71, 1.4%), with all other countries presenting frequencies below 1%. In East Africa, a single case (1/1907, 0.5%) was observed in Kenya and this dated back to 1998. In Southeast Asia, *pfcrt* amplifications were observed in Laos (n=9, 0.55%) and Vietnam (n=3, 0.15%). Five different breakpoint ranges were identified across the amplifications, ranging in size from ∼23 to 59 Kb: all shorter than the ∼100 kb duplication observed in Sabah (Table 3). In all cases, the breakpoints were in either polyA and polyT tracts. In addition, two samples from Ghana had amplifications of approximately 20 kb for which breakpoints could not be determined but are likely distinct from the other breakpoints identified. In Ghana, Gambia and Burkina Faso, four, two and two different breakpoints were observed respectively. The most common amplification in Ghana, and across the 86 non-Sabah cases, was one of the shortest amplifications observed, spanning 22.87 kb. The earliest observed amplification was the Kenyan case from 1998, with all other cases observed between 2007 and 2020 (Figure 3b). The highest frequency was observed in 2008, though there is a large amount of variation between years. As the Pf8 dataset aggregates samples across multiple independent studies collected opportunistically at different locations at different times, rather than according to a unified sampling framework, these frequencies should be interpreted with caution and may not reflect the true prevalence of pfcrt amplification across these regions.

**Figure 3.**
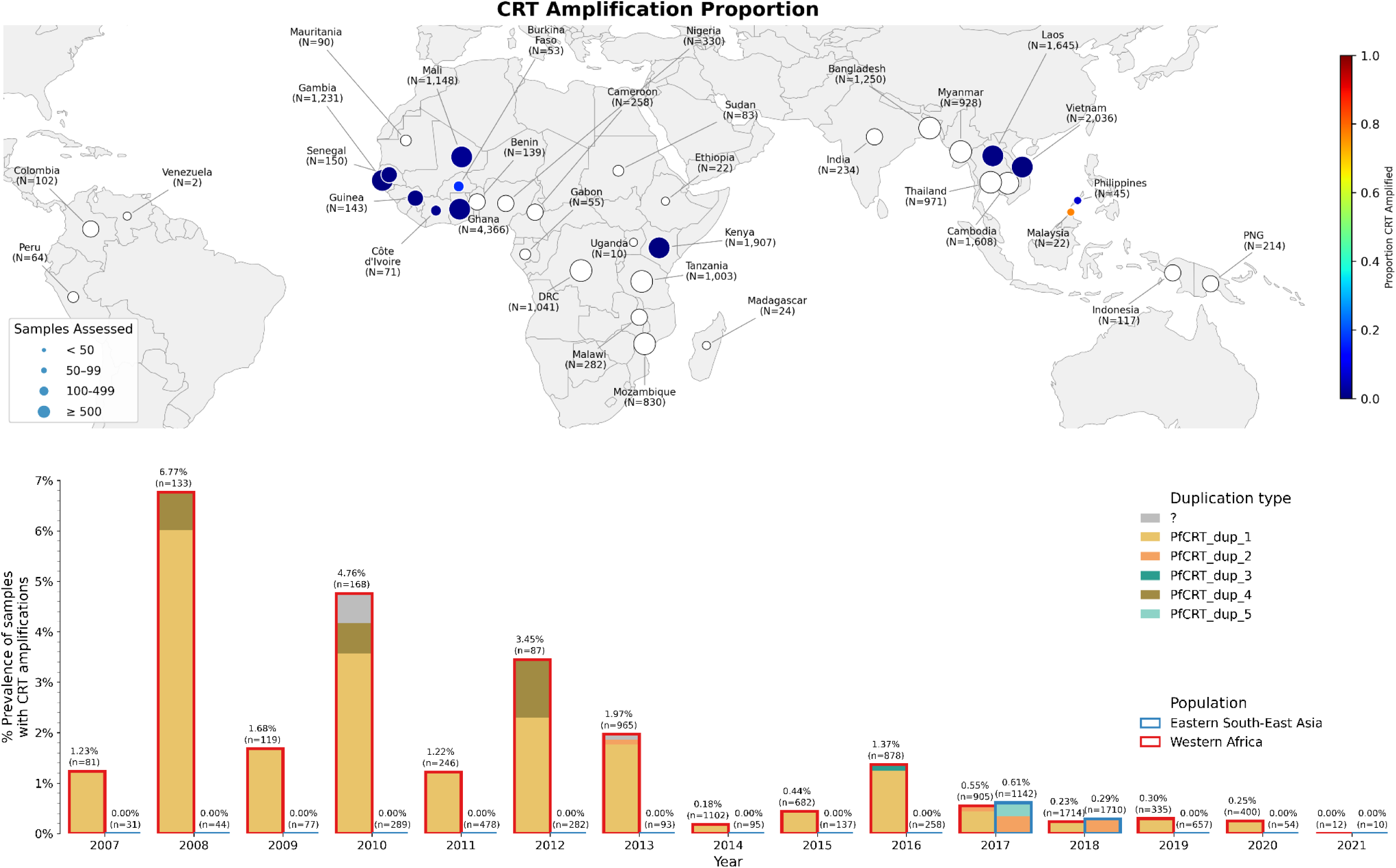
Global spatial and temporal distribution of *pfcrt* amplifications. Panel (a) illustrates the spatial distribution of *pfcrt* amplifications detected in samples with non-missing *pfcrt* CNV calls from the MalariaGEN Pf8 data release and using qPCR data for cases from Malaysia and the Philippines. Countries where no samples with *pfcrt* amplifications were detected are shown with white circles. Panel (b) presents the yearly prevalence of *pfcrt* amplifications in the MalariaGEN Pf8 data release since the first year amplifications were detected. Above each bar is the % prevalence of CRT amplifications in samples with CNV calls in CRT, and the total number of samples with non-missing CNV calls in CRT are shown as n. One Kenyan sample with *pfcrt* amplification presented in 1998 is not shown. The Pf8 dataset is a multi-study compilation in which samples were collected opportunistically rather than according to a unified sampling framework, and may therefore not reflect the true distribution of parasite populations across space or time.

**Table 3.** Geographical distribution of samples outside of Malaysia and the Philippines with evidence of *pfcrt* amplification. Breakpoints were taken from the MalariaGEN Pf8 release and, where the data were missing, were resolved manually using IGV. For the two samples from Ghana whose breakpoint is “Uncertain”, we were unable to identify precise breakpoints based on faceaway read pairs, and instead approximated the size of the amplification based on data from the coverage-based HMM method.

| CRT breakpoint | Breakpoint range | Amplification length | Country | Frequency |
| --- | --- | --- | --- | --- |
| PfCRT_dup_1 | 398,869~398,898 - 421,765~421,808 | 22,867~ | Ghana | 0.8% (33/4366) |
|  |  |  | Mali | 1.7% (20/1148) |
|  |  |  | Burkina Faso | 15% (8/53) |
|  |  |  | Côte d'Ivoire | 1.4% (1/71) |
|  |  |  | Gambia | 0.08% (1/1230) |
|  |  |  | Guinea | 0.7% (1/143) |
|  |  |  | Senegal | 0.7% (1/150) |
| PfCRT_dup_2 | 399,263~399,291 - 422,103~422,136 | 22,812~ | Laos | 0.5% (9/1645) |
|  |  |  | Ghana | 0.05% (2/4366) |
| PfCRT_dup_3 | 390,397~390,416 - 421,765~421,808 | 31,349~ | Gambia | 0.08% (1/1230) |
|  |  |  | Kenya | 0.05% (1/1907) |
| PfCRT_dup_4 | 393,103~393,133 - 421,765~421,808 | 28,632~ | Ghana | 0.05% (2/4366) |
|  |  |  | Burkina Faso | 1.9% (1/53) |
| PfCRT_dup_5 | 365,968~365,991 - 425,216~425,236 | 59,225~ | Vietnam | 0.1% (3/2036) |
| Uncertain | 399,001 ~ 419,000 | 20,000~ | Ghana | 0.05% (2/4366) |

### Distinct origin of pfcrt duplications from different countries

Exploration of the sequence composition of the cases with evidence of *pfcrt* duplication was conducted to assess the similarity of the haplotypes within each case using samples from the MalariaGEN Pf8 data release. Amongst 47 clonal cases (Fws ≥ 0.99) with high-quality data (non-missing CRT mutation calls) within the *pfcrt* gene, 94% (44/47) had heterozygote calls at 3 or more loci indicative of heterologous haplotypes within the duplication (Figure 4). The heterologous haplotypes all comprised combinations of 76K (wild-type) and 76T (mutant) variants. Three cases carried homozygote calls at all *pfcrt* codons assessed, with mutant 76T genotypes. Samples from the Sabah outbreak contained heterozygous mutations in *pfcrt* not seen elsewhere (L160Y, I166V, H273N, N326D) suggesting a distinct origin of this duplication (Figure 4). Outside of the Sabah outbreak, in samples with duplications of *pfcrt* there was a high degree of genetic relatedness between eight of nine cases from Laos, between the two cases from Vietnam, and between two of three cases from Burkina Faso. These highly related samples were the exception and most genetic distances between samples with *pfcrt* amplifications were similar to genetic distances between any pair of randomly selected samples in Pf8, despite the amplifications having identical breakpoints and therefore likely being inherited from the same initial amplification event. This suggests that these amplifications in *pfcrt* can persist in parasites and be inherited stably across several generations, and are not just seen transiently as part of clonal expansions.

**Figure 4.**
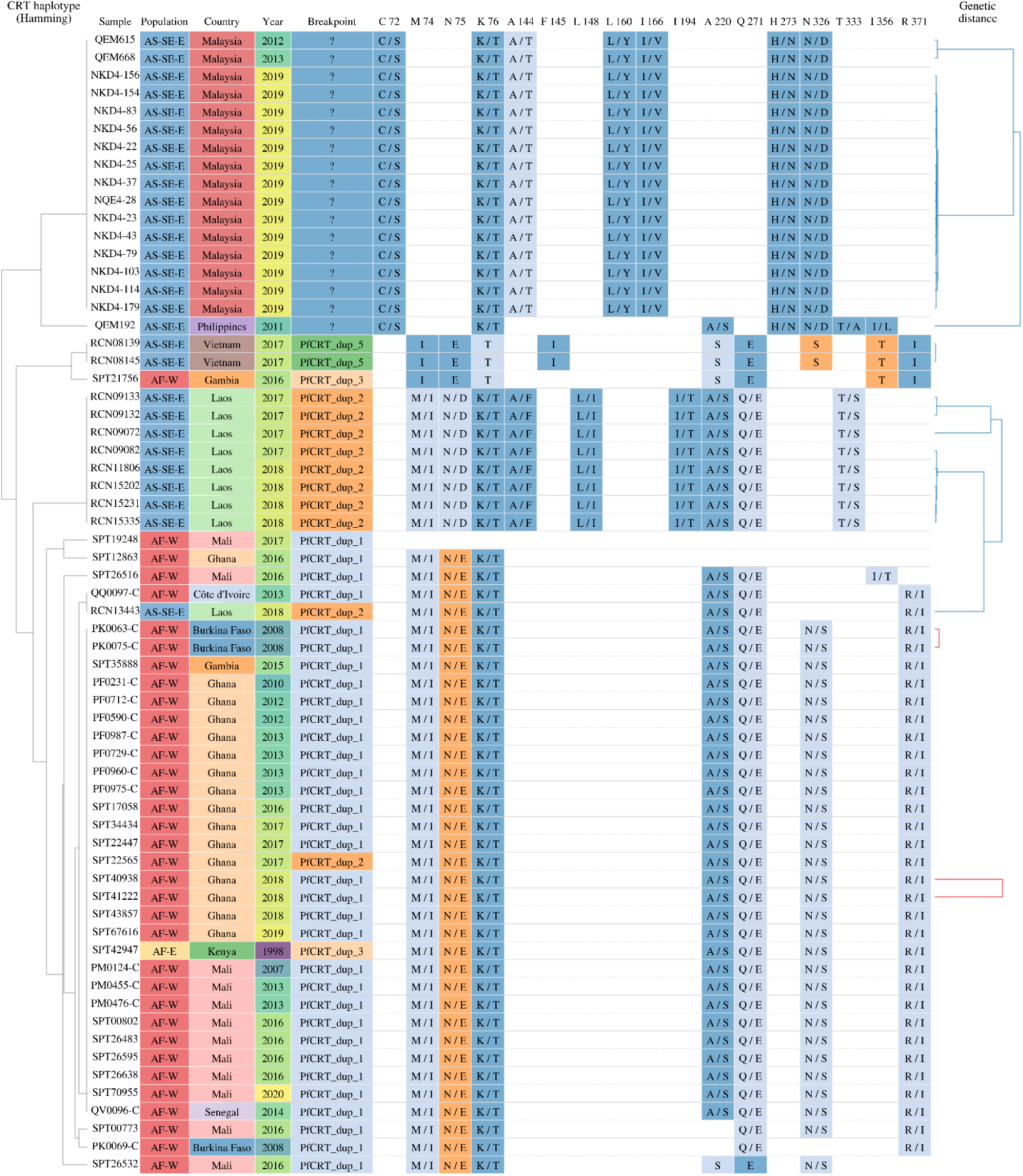
Graphic summary of *pfcrt* variants and relatedness amongst global samples with *pfcrt* amplification. The data was derived from whole genome sequencing calls in monoclonal samples from MalariaGEN Pf8 with resolved amplification breakpoints, or Malaysia with evidence of *pfcrt* amplification. In MalariaGEN Pf8, evidence of amplification was derived from genomic signals, whilst in Malaysia, evidence was derived from qPCR data. For each sample, the following are shown: country and population of origin, CNV breakpoint classification and amino acid changes across *pfcrt* (homozygous amino acid changes, heterozygous amino acid changes, or no change (in blank) — there are no missing positions). Where heterozygous, the REF amino acid is shown first, followed by the ALT amino acid. To the left is a tree constructed using the het SNP pattern displayed in the table, and to the right is a tree constructed using whole-genome genetic distance (where red branches connect African samples and blue branches connect Asian samples, and distant branches are hidden for visual clarity).

## Discussion

Our investigation of a *P. falciparum* outbreak in Sabah Malaysia identified a novel 100 Kb amplification encompassing the *pfcrt* locus. Whilst Malaysia has recently eliminated *P. falciparum*, our exploration of the open-access MalariaGEN repository comprising data from endemic countries across the globe revealed *pfcrt* amplifications in 11 other countries, with a notable frequency in West Africa. Although point mutations in *pfcrt* have been implicated in resistance to chloroquine (CQ), piperaquine (PPQ) and amodiaquine (AQ), the impact of gene amplification on drug resistance has not been explored in depth. A previous study detected bioinformatic signatures of *pfcrt* amplifications in an earlier MalariaGEN data release but these were not confirmed with experimental data and thus remained putative (Ravenhall et al., 2019). Our investigations in Sabah and the Philippines provide robust confirmation of *pfcrt* amplification as a polymorphism circulating in natural parasite populations. We discuss the dynamics of the Sabah outbreak and examine country-wide genomic data to evaluate the potential adaptive role of *pfcrt* amplification in response to antimalarial drug pressure.

The *pfcrt* amplification was first detected at high frequency amongst the 2019 Banggi island outbreak cases in Sabah. The clonal dynamics of the outbreak infer that a single strain entered the population and expanded rapidly, consistent with an imported index case. The predominance of the 100 Kb amplification amongst the outbreak cases infers a potential role as an adaptive driver. No other validated or candidate drug resistance mutations were detected that might explain a selective expansion. There are 24 other genes in the 100 kb amplification region but *pfcrt* is a compelling candidate given its known and extensive drug resistance history. However, it is possible that the Banggi outbreak was not driven by an adaptive force, but rather reflected neutral epidemiological processes resulting from the introduction of a new parasite strain into a human population with relatively low immunity. Indeed, several clonal expansions of *P. falciparum* were observed in Sabah in the years preceding elimination, with no clear evidence of resistance drivers. Clonal expansions of *P. vivax* were also observed in Sabah in the pre-elimination phase, with enrichment of several potential resistance markers, but resistance mechanisms for this species are less well understood than for *P. falciparum* (Auburn et al., 2018).

The origin of the Sabah *pfcrt* amplification is also unclear. Malaysia experiences frequent malaria importation, with neighbour-joining analyses showing close genetic identity between cases from Sabah and the Philippines. Shared mutant *pfcrt* haplotype profiles were observed between Sabah and Palawan island, suggesting that parasite movement between the countries may have been historically frequent. The region appears to have harboured *pfcrt* amplifications since as early as 2011, with earlier years not assessed here. Data from the open-access MalariaGEN repository identified a single Kenyan case with amplification dating back to 1998, followed by multiple amplifications in west Africa since 2007, although this pattern may be influenced by the temporal span of the available data. The relationship between the Malaysian/Philippines and West African amplifications is not clear but these are likely to reflect distinct origins given the distinct breakpoints and generally high mutation rate of amplifications (Nair et al., 2007).

Several lines of evidence suggest an adaptive role of *pfcrt* amplification. Most notably, the extensive size of the amplification (up to 100 kb) is suggestive of an adaptive benefit to the parasite that would need to outweigh the fitness cost of generating this extra DNA and presumably associated protein. Indeed, ex vivo studies of *pfmdr1* amplification indicate decreased survival fitness in the absence of drug pressure (Preechapornkul et al., 2009). Furthermore, the frequency of resistance-associated parasite amplifications has been observed to drop rapidly after drug removal (Auburn et al., 2018; Preechapornkul et al., 2009).

Evidence of multiple breakpoints across the global dataset may indicate that *pfcrt* amplification has arisen on several independent occasions, supporting an adaptive role. Six different breakpoint ranges were observed in the Pf8 dataset (Table 3), and the Sabah outbreak samples constitute a seventh unique breakpoint, suggesting there have been at least seven independent origins. Indeed, in Ghana alone four different breakpoints were observed. Whilst resistance-associated point mutations in genes such as *pfcrt* and *pfdhfr* demonstrated global spread from a handful of origins, multiple amplifications of genes such as *pfmdr1* have been observed in a single population owing to the relatively higher mutation rate (Nair et al., 2007). In the majority of *pfcrt*-amplified cases, the presence of a wild type (3d7) *pfcrt* copy alongside a highly mutated copy, whose mutation profile reflects locally circulating haplotypes, is most parsimoniously explained by sexual recombination and crossing over than by tandem duplication followed by independent re-evolution of an existing local haplotype. Two recombination-based mechanisms are plausible. One possibility is unequal crossover in a single event during meiosis, where chromosomes from two parasites carrying different *pfcrt* haplotypes misalign at low-complexity flanking sequence and cross over at non-allelic positions, simultaneously generating the duplication and acquiring the diverged copy. An alternative possibility is tandem duplication within a single lineage followed by interlocus gene conversion or crossing over in a subsequent sexual recombination event that replaces one copy with a diverged haplotype from a co-infecting parasite. Both the identification of faceaway read pairs near breakpoints and the existence of parasites carrying homozygous *pfcrt* amplifications (in Vietnam and the Gambia) lend validity to the latter mechanism, demonstrating that tandem duplication is a stable, heritable event that can become established in parasite populations independently of acquiring a diverged copy. Consistent with either mechanism, breakpoints identified in these samples fall in polyA and polyT tracts, which are well-established hotspots for non-allelic homologous recombination due to misalignment during meiosis. The inability to detect faceaway reads in some samples may reflect the fact that Illumina sequencing technology has difficulty base-calling after sequencing through long homopolymer repeats.

If *pfcrt* amplification is assumed to have an adaptive benefit mediated by drug resistance, critical follow-up questions are which drug(s) is impacted and by which mechanism(s). A range of antimalarial drugs are advised as first-line policy for *P. falciparum*, or co-endemic *Plasmodium* species, in the countries where amplifications were observed. Aside from the Sabah findings, the highest frequency of amplifications was observed in West Africa, with only a single East African case in 1998. East and West Africa have historical differences in antimalarial drug use including recommendations for artemisinin-based combination therapies (ACTs) (Walker et al., 2025). In West Africa, artesunate plus amodiaquine (ASAQ) and artemether-lumefantrine (AL) are prescribed for uncomplicated *P. falciparum* infection, whilst countries in East Africa tend to use AL only. The difference in amodiaquine (AQ) drug pressure between East and West Africa could potentially explain the difference in *pfcrt* amplification frequency. To date there has been limited evidence of therapeutic efficacy failures against artesunate-amodiaquine in West Africa but effective artemisinin therapy could conceal any partner drug failures (WHO, 2025b). Only sporadic cases of *pfkelch13* mutations associated with artemisinin resistance have been detected in west Africa, suggesting efficacy remains high (WHO, 2025b). There are no validated markers of AQ resistance, only two candidates in *pfcrt* and *pfmdr1* based on laboratory and epidemiological data (WHO, 2025a). While a role in AQ resistance is compelling, other antimalarials used in the affected populations cannot be excluded as selective determinants of *pfcrt* amplification. Indeed, in several of the countries where both ASAQ and AL are prescribed, the latter therapy is more commonly used and hence likely to impart greater selective pressure from lumefantrine (LF) than AQ (WHO, 2025b). The Illumina data on monoclonal *pfcrt*-amplified cases from the MalariaGEN Pf8 repository revealed that the majority of cases carried both wild type and mutant variants at amino acid 76, associated with chloroquine resistance. Heterozygotes were also observed at a range of other drug resistance-associated *pfcrt* variants, but phasing of the haplotypes was not readily permitted with short-read Illumina data. Long-read sequencing of *pfcrt* in Sabah and the Philippines was highly insightful, revealing a complement of wild type (3d7) and mutant *pfcrt* haplotypes in approximate 1:1 ratios indicative of 1 copy of each haplotype. The mutant haplotype was most similar to the previously described PH2 haplotype from the Philippines, which includes mutations 72S, 76T, 144T, 160Y and 326D (Chen et al., 2003; Petersen et al., 2015). An additional mutation, 166V, was observed on the PH2 background in the mutant haplotypes within the amplification. Although the resistance profile of the 166V mutation is unclear, laboratory tests using genetically engineered parasite clones carrying a variety of *pfcrt* haplotypes provides useful insights on the susceptibility of the PH2 haplotype to a range of antimalarial drugs (Chen et al., 2003; Petersen et al., 2015). These experiments indicate that the PH2 haplotype confers a 4.7-fold increase in IC50 to CQ (53-54 nM) relative to the wild-type haplotype, at the cost of reduced growth in vitro. The PH2 haplotype also confers a 4-fold increase in IC50 to monodesethyl-amodiaquine (45 nM), a clinically relevant metabolite of AQ. In contrast, the PH2 haplotype has significantly increased susceptibility to LF and quinine relative to the wild-type haplotype, but no difference with piperaquine.

Several drugs including AQ and CQ are thought to act upon the parasite food vacuole by interfering with haem detoxification, leading to the accumulation of toxic haem and subsequent parasite death (Gorka et al., 2013). In vitro investigation using CQ indicates that the PH2 haplotype reduces accumulation of drug in the digestive vacuole (Chen et al., 2003; Petersen et al., 2015). However, the same study identified reduced accumulation of CQ across a variety of haplotypes with varying CQ IC50s suggesting that additional mechanisms of resistance may be at play for this and potentially related drugs.

Whilst a wealth of information is available on the effect of single *pfcrt* haplotypes on susceptibility to a range of drugs, the combined impact of multiple haplotypes has not been investigated. With potentially opposing effects, the susceptibility and fitness consequences of combined mutant and wild-type haplotypes is particularly complex to predict. In theory, the wild-type haplotype could potentially override the growth cost observed in mutant PH2 haplotypes, producing a more adaptive genetic background. Functional studies using schizont maturation assays in natural infections or genetically engineered wild-type and mutant *pfcrt* haplotype combinations are needed to confirm drug susceptibility, growth and other mechanistic profiles for a range of antimalarials. Further studies are also needed to determine the complement of haplotypes observed in different geographic regions and their potential drug modulatory effects.

Other research gaps that need to be addressed include assessment of *pfcrt* expression and translation to confirm that the amplifications lead to increased RNA and protein. Owing to restricted sample material in Malaysia and the Philippines, and the use of open-access data from the MalariaGEN dataset, this was not directly assessed in our studies. However, the architecture of the amplified sequence region appears conducive to functional expression of full copies of both *pfcrt* haplotypes.

Although we detected *pfcrt* amplifications in a wide range of malaria-endemic countries, our frequency approximations are likely to be an underestimate owing to constraints in detecting amplifications using short-read sequence data in polyclonal infections and data derived from sWGA-amplified material. Wider surveillance is needed using targeted assays such as the qPCR-based TaqMan assay developed in this study. As such, this study suggests the need for a scheduled malaria screening programme along the Malaysia- Philippine borders. This strategy is important to prevent malaria re- establishment in Malaysia. Certain sentinel stations that conduct malaria screenings along these borders would provide better and justified evidence to invest on molecular and genomic screening for malaria; such as the *pfcrt* amplification technique.

In conclusion, our study reveals potentially important new variants of *pfcrt* including one of the largest amplifications (100 Kb) reported for *P. falciparum*. These previously underreported *pfcrt* amplifications appear to be circulating in sites across the globe including highly endemic regions of west Africa, with potentially critical implications for antimalarial treatment efficacy. In a broader evolutionary context, our discoveries highlight the remarkable plasticity of *Plasmodium* and the critical need to maintain genomic discovery within local surveillance programs.

## Methods

### Master table of all samples used in the study

A master table listing all samples used in the study is provided in Supplementary Data 1. The master table indicates which analyses each sample was used in and where the genetic data was derived from. Further details on individual studies are provided below.

### Patient sampling and demographics in Pulau Banggi outbreak

*P. falciparum* cases were identified on Banggi Island, Kudat District, Sabah, Malaysia between January and June 2019 through three surveillance modalities implemented by the Sabah State Health Department: mass blood surveys (MBS), active case detection of close contacts of identified cases (ACD), and routine passive health facility case detection (PCD). Demographic information, including age, sex, occupation and village of residence were recorded (Supplementary Table 4). Cases were classified as imported or introduced cases through Malaysian public health investigation reporting, with recorded recent residence or travel, and attributed exposure to the neighbouring island of Balabac (∼70km away) in Palawan Province, Philippines. Venous blood was collected from 102 cases as part of the mandatory PCR confirmation of *Plasmodium* species at the State Public Health Laboratory, of which 98 had sufficient volume and parasite gDNA able to be obtained for additional *pfcrt* sequencing.

### Patient sampling within the Sabah state-wide survey

From 2010 to 2015, non-pregnant patients with microscopy positive malaria were enrolled as part of a prospective malaria pathophysiology study (Malaysian MREC ethics approval: NMRR-10-754-6684) after presenting to either an adult tertiary hospital site (≥12 years of age), or 3 district hospital sites (Kudat, Kota Marudu and Pitas; any age) in Sabah, Malaysia, with written informed consent obtained (Barber et al., 2013; Grigg, William, Barber, et al., 2018; Grigg, William, Piera, et al., 2018). Demographic, clinical, and epidemiological data were obtained, and PCR was performed for confirmation of *P. falciparum* and other species.

From 2016, patients presenting with fever of any age from routine public health microscopy positive malaria notifications from across Sabah (encompassing 24 district hospitals) were included as part of a state-wide malaria surveillance study (NMRR-15-168-24821), with PCR results obtained from the State Public Health laboratory, and additional clinical and epidemiological data collected by case record forms including assessment of WHO severe malaria criteria.

### Patient sampling in the Philippines

Patient samples from comparator sites in the Philippines were collected by the Department of Parasitology-Malaria National Reference Laboratory of the Research Institute for Tropical Medicine, through regular malaria-related activities: (a) therapeutic efficacy studies (TES)and (b) National External Quality Assessment Scheme (NEQAS). Archived samples used in this study were microscopy-confirmed *P. falciparum* and were collected from consenting participants prior to enrollment or blood collection.

TES samples were obtained from eligible participants aged 6 months to 59 years old through health-facility or village-based surveys in selected malaria-endemic sites in Palawan and across several periods: 2013 (Bataraza, Brooke’s Point and Rizal); 2015, 2017-2018 (Bataraza and Rizal); and 2025 (Rizal), the most recent from drug efficacy surveillance integrated into routine case detection.

NEQAS samples were collected from 2013 to 2026 from patients aged 12 years and older who tested positive for malaria during routine screening at health facilities in Rizal, Palawan. Following sample collection, patients from both the TES and NEQAS were referred for clinical management and treatment in accordance with national malaria guidelines.

### Open-access MalariaGEN comparator data

Published *P. falciparum* Illumina whole genome sequencing data was derived from the MalariaGEN Pf8 community project release (Malaria Genomic Epidemiology Network (MalariaGEN) et al., 2025). The data comprises 24,409 high quality samples from 34 countries across the globe and derives from both gDNA and sWGA libraries sequenced on a range of Illumina platforms to generate mostly 150bp paired-end reads (some earlier samples were sequenced with shorter reads). A total of 22,047 (90% from the high quality samples) of the MalariaGEN Pf8 cases were investigated for evidence of *pfcrt* copy number amplification. These cases were selected based on those without missing CNV calls.

A subset of 300 (1.2% out of 24,409) MalariaGEN Pf8 comparator samples were included in geo-mapping analyses to determine the likely provenance of the Banggi outbreak cases. The sample subset was selected to achieve representation of each of the global regions defined in the MalariaGEN Pf8 report based on combined epidemiological and genetic clustering information; South America (SA), Western Africa (AF-W), Central Africa (AF-C), Northeastern Africa (AF-NE), Eastern Africa (AF-E), Eastern South Asia (AS-S-E), Far East South Asia (AS-S-FE), Western Southeast Asia (AS-SE-W), Eastern Southeast Asia (AS-SE-E) and Oceania-New Guinea (OC-NG). A random selection of 30 monoclonal cases (defined as cases with Fws>0.95 as documented within the MalariaGEN Pf8 report) were selected from the high-quality sample set for each region.

### DNA extraction and species confirmation

For the samples from the Pulau Banggi and Sabah state-wide survey, DNA extraction was undertaken on ∼200 ul whole blood extracts using commercial kits (QIAamp mini blood kit, Qiagen), and *Plasmodium* species was confirmed using a real-time PCR assay (Rougemont et al., 2004). For the samples from the Philippines, DNA extraction was undertaken on dried blood spots using commercial kits (QIAamp DNA mini kit, Qiagen) and *Plasmodium* species was confirmed using nested PCR (Calderaro et al., 2007; Lee et al., 2011; Snounou & Singh, 2002). Aliquots of each sample were subject to selective Whole Genome Amplification (sWGA) for *P. falciparum* using a previously described protocol (Oyola et al., 2014).

### Illumina amplicon-based sequencing and variant calling

Illumina amplicon-based sequencing was undertaken on sWGA product using the SpotMalaria *P. falciparum* genetic barcode, which comprises drug resistance targets, 100 neutral epidemiological markers and 2 mitochondrial *Plasmodium* species confirmation targets (Jacob et al., 2021). The drug resistance targets include validated SNP-based resistance markers in *pfcrt*, *pfkelch13*, *pfmdr1*, *pfdhfr* and *pfdhps*. Copy number variation cannot be robustly detected by amplicon-based sequencing and hence are not incorporated in the assay. Library preparation was undertaken using a previously described protocol (Jacob et al., 2021). Briefly, target amplification was conducted in three separate multiplex PCR reactions, followed by incorporation of unique indexes for each sample. The resulting libraries were subject to sequencing on a MiSeq platform using 300 cycles generating 150bp paired end reads.

Variant calling was performed using targeted variant caller workflow implemented in the vivaxGEN NGS-Pipeline (https://github.com/vivaxgen/ngs-pipeline) using the Pf3D7_v3 configuration (https://github.com/vivaxgen/vgnpc-plasmodium-spp/tree/main/Pfalciparum/Pf3D7_v3). Briefly, raw reads were mapped to the Pf3D7_v3 reference genome using minimap2 (Li, 2021). Targeted genotyping for the 100 SPOTMalaria barcode SNP positions was conducted using freeBayes to generate individual VCF files for each sample (Garrison & Marth, 2012). A final multisample VCF file was generated by merging individual outputs. Genotypes were then filtered based on a minimum total depth of 10 and heterozygote calls were defined by a minor allele read depth > 5 and a minor allele read ratio > 0.1.

To assess sequencing data for drug resistance associated variants, the unfiltered multisample VCF was subset to include regions containing WHO validated genes associated with drug resistance. This was done using the *malaria-popgen-toolkit* missense-drugres-af command (https://github.com/aosborne13/malaria-popgen-toolkit). The regions included: *pfcrt* (Pf3D7_07_v3: 402385 - 406341), *pfmdr1* (Pf3D7_05_v3: 955955 - 963095), *pfdhps* (Pf3D7_08_v3: 547896 - 551057), *pfdhfr* (Pf3D7_04_v3: 747897 - 750065), and *pfk13* (Pf3D7_13_v3: 1724600 - 1727877). Variants within the subset VCF files were annotated using the *BCFtools* consequence calling command to determine the corresponding amino acid shift or variant type. Only variant positions within samples containing a read depth (DP) ≥ 5 were included in allele frequency estimates and sample proportion estimates. Positions with more than one alternative allele were retained if a DP ≥ 5 was retained. The variant proportion within the population was calculated from the number of samples containing the alternate allele out of the total number of samples. The number of heterozygous samples within a population was calculated as the number of samples containing the heterozygous genotype call (0/1).

### Nanopore amplicon-based sequencing and variant calling

Single-molecule nanopore sequencing of the full length *pfcrt* gene was undertaken either using genomic DNA or sWGA product as an input for the amplification of the gene. In brief, a 3331 bp amplicon spanning Pf3D7_07_v3:403136-406466 was generated using a primer pair designed by David A. Fidock’s lab (Columbia University, USA; *unpublished*) as shown in Supplementary Table 4 with the following PCR reactions which was done in Axygen 96 well PCR microplate (Corning Life Sciences, Corning, USA) in 50 µL, containing 10 µL of 5X PrimeSTAR GXL Buffer (TakaraBio, Japan), 200 μM dNTP mixture, 300 nm of each forward and reverse primer, and 2 µL of template DNA. PCR conditions used: an initial denaturation at 98 °C for 30 seconds, followed by 30 cycles of denaturation at 98 °C for 10 seconds, annealing at 54 °C for 15 seconds and extension at 68 °C for 3.5 minutes. A final extension at 68 °C for 4 minutes, and the reaction was held at 4 °C until removed from the thermocycler. PCR amplicons were purified using the QIAquick PCR purification kit (Qiagen, Germany) according to the manufacturer’s instructions and quantified using Qubit™ dsDNA HS Assay Kit (Invitrogen, USA). Amplicon sequencing was performed using Oxford Nanopore Technologies’ Barcoding Kit 96 (SQK-NBD114.96, Oxford Nanopore Technologies, Oxford, UK) following the manufacturer’s instructions. Libraries were prepared from PCR amplicons and sequenced on a MinION Mk1C using a R10.4.1 flow cell (FLO-MIN114). Basecalling was performed using the super-accuracy (SUP) method employed by dorado version 1.3.1. Long haplotypes were resolved using the vivaxGEN NGS-Pipeline long haplotype workflow, which employed a two-stage discovery and targeted genotyping workflow. Briefly, long reads were mapped to the *P. falciparum* 3D7 (Pf3D7_v3) reference genome using minimap2. De novo variant calling was performed on individual samples with Clair3 to generate GVCFs (Zheng et al., 2022), which were subsequently joint-called into a multi-sample VCF using GLNexus (Yun et al., 2021). To ensure complete allele information for all samples to generate long haplotypes, these discovered coordinates were merged with a predefined panel of SNPs-of-interest. Clair3 was then employed for targeted genotyping at these combined positions across all individual BAM files, ensuring genotype recovery for both variant and invariant sites. Read-level haplotypes were reconstructed using a custom Python script to extract alleles from aligned BAM reads at the designated VCF coordinates. Long haplotypes with read depth ratio over total reads < 0.05 were discarded. Samples with <100X read coverage were marked as low quality samples.

### Quantification of Pfcrt copy number by TaqMan qPCR

*Pfcrt* copy number was assessed by TaqMan quantitative PCR (Applied Biosystems QuantStudio™ 6 Flex Real-Time PCR System; Thermo Fisher, USA). Primers and probes for *Pfcrt* and *β-tubulin* (single copy reference) were designed by the Integrated DNA Technologies designing team and are listed in Supplementary Table 4. The *Pfcrt* probe was labelled with reporter dye 6-carboxyfluorescein (FAM) and the 3′-quencher BHQ1. The *β-tubulin* probe was labelled with reporter dye Cyanine5 (Cy5) and the 3’-quencher BHQ2. TaqMan qPCR reactions were done as multiplex in MicroAmp 96-well plates (Applied Biosystems,Thermo Fisher Scientific, USA) in 20 µL, containing 10 µL of TaqMan™ Fast Advanced Master Mix for qPCR (Thermo Fisher, USA), 400 nm of each forward and reverse primer, 200 nm of each probe and 2 µL of template DNA. The qPCR program was as follows: UNG activation at 50 °C for 2 minutes, polymerase activation at 95 °C for 20 seconds, and 45 amplification cycles of 95 °C for 3 seconds and 60 °C for 30 seconds. Relative copy number was calculated using the 2^−ΔΔCt^ method. 3D7 was used as the control sample. All samples were analysed in duplicate and were excluded if the Ct values were >35.

### Illumina whole genome sequencing and variant calling for CNV analysis

Illumina whole genome sequencing (WGS) of the Banggi outbreak and state-wide survey samples from Malaysia was conducted at the Wellcome Sanger Institute, Cambridge, within a MalariaGEN *P. falciparum* community project. Samples that demonstrated successful sequencing at the SpotMalaria barcode were subject to library preparation using the standard Illumina protocol. All libraries were prepared using sWGA material. Sequencing was conducted on an Illumina platform using 300 cycles to generate 150 bp paired end reads.

Samples processed by Illumina WGS were initially genotyped using the MalariaGEN Pf7.x pipeline (coding SNPs variant calling pipeline) for Supplementary Figure 3B, before reprocessing using the MalariaGEN Pf8 snp-indel-calling pipeline (coding and non-coding SNPs and indels, https://github.com/malariagen/malariagen-pf8-snp-indel-calling) for higher resolution analysis (Supplementary Figure 3A). The BAM files from the latter were then analysed using the Pf8 cnv-calling pipeline (https://github.com/malariagen/malariagen-pf8-cnv-calling), trained against samples from the Pf8 dataset. Determination of breakpoints involved identifying faceaway read pairs on IGV for evidence of tandem duplications.

### Population genetic analysis

For population genetics analysis, the raw reads from Malaysian samples and Pf8 comparator samples were processed using the GATK-based discovery variant caller workflow implemented in the vivaxGEN NGS Pipeline (https://github.com/vivaxgen/ngs-pipeline) with Pf3D7_v3 configuration (https://github.com/vivaxgen/vgnpc-plasmodium-spp/tree/main/Pfalciparum/Pf3D7_v3). Briefly, the reads were mapped to both Pf3D7_v3 and human reference sequences using bwa-mem2. Read pairs were then filtered out from the map file when any of the pairs read mapped to the human reference genome. The map files were then mate-fixed and deduplicated to generate the final map file. Base recalibration was performed on the final map file based on the high quality SNP list from Pf7 dataset using GATK. Individual variant calling was then performed using GATK to generate GVCF files. The GVCF files of each sample were then jointly genotyped using GATK GenotypeGVCFs to produce the final multisample VCF files. GATK version 4.6 was used for all these steps.

Population genetic analysis was performed using the MEA-Pipeline (https://github.com/vivaxgen/mea-pipeline) on the WGS data. The variants were filtered with query notation of: `initial-variants-HETd2r0.1-d5-MAC3-V0.5-split-dedup-MAF0.05-S0.25-V0.25-MAF0.1`, resulting in 116,704 biallelic SNPs from 634 samples (333 Sabah, 301 non-Sabah). A neighbor-joining tree was constructed from a proportional genetic distance matrix generated using Ape software (Popescu et al., 2012). The Fws score for each sample was calculated using moimix software (https://bahlolab.github.io/moimix/) with monoclonal samples defined as those with Fws > 0.95. Pairs of samples were considered clonal when their genetic identity was greater than 99.9% (less than 117 base differences out of 116,704 SNPs).

For population genetic analyses based on the amplicon sequencing data, the 100 SNP diversity loci included in the SPOTMal panel were extracted from the WGS dataset and combined with the available SPOTMal amplicon sequencing data from Sabah and the Philippines, resulted in total of 690 samples (350 Malaysian, 40 Philippines and 300 others). For samples having both WGS and amplicon sequencing data, the data set with fewer missing genotypes was selected for further analysis. A neighbor-joining tree was then constructed using the same approach as for the WGS dataset, employing the Ape package on a proportional genetic distance matrix.

## Acknowledgements

Sample collection and sample processing were supported by the Ministry of Health, Malaysia (grant number BP00500420 and grant number BP00500/117/1002) to GSR; the Australian National Health and Medical Research Council (grant numbers 496600, 1037304 and 1045156); the US National Institutes of Health (grant numbers R01AI116472-03 and 1R01AI160457-01) to TW and GSR, and the UK Medical Research Council, Natural Environment Research Council, Economic and Social Research Council, and Biotechnology and Biosciences Research Council (grant number G1100796). Analysis was supported by the National Health and Medical Research Council of Australia (PP2001083 awarded to S.A., fellowships to NMA [1042072], MJG [1138860] and BEB [2016801, 2016792]. IDSKKS and Menzies research staff are supported through the RESPOND project, Indo-Pacific Centre for Health Security, Department of Foreign Affairs and Trade, Australian Government. The whole genome sequencing component of the study was supported by the Medical Research Council and UK Department for International Development (award number M006212 to D.K.) and the Wellcome Trust (award numbers 206194 and 204911 to D.K.). TaqMan assay and ONT sequencing assay were supported by the Australian Centre of Research Excellence in Malaria Elimination (awarded to A.R.). Sample collection and processing for TES and NEQAS were supported by the Department of Health and RITM-NRL, with additional support in 2024 from the Joint WPRO/TDR Impact Grants (grant number AP23-01266 MAL PHL). We thank the patients who contributed their samples to the study, and the health workers and field teams who assisted with the sample collections. Whole genome sequencing was undertaken by the Wellcome Sanger Institute. We thank the staff of the Wellcome Sanger Institute for contributions to samples logistics, sequencing and informatics. We thank the Director-General, Ministry of Health, Malaysia, for permission to publish this manuscript.

## Author Contributions

S.A., M.J.G., N.M.A., G.S.R and B.E.B. conceived the study. S.A., R.P., and M.J.G. designed the study. D.P.K and R.P. contributed to whole genome sequencing data production. A.R., H.T., J.H.A.L, M.L.M., D.T, J.L., K.P., D.F. and S.A. contributed to the laboratory work to generate qPCR and Oxford Nanopore Technologies sequencing data. C.L., H.T., A.O., K.S.H., J.W., and R.P. contributed to bioinformatic pipeline development and data analysis. G.R., J.H.A.L., M.L.M., J.L., D.T., S.G., A.J., A.M.M., W.G., F. E.E. and T.M. contributed essential field-based malaria collections and metadata. S.A., A.R., C.L., H.T., and A.O. wrote the original manuscript draft. All authors provided guidance on the study design and interpretation and read and approved the final manuscript.

