## Supplementary figures for "*Pfcrt* copy number amplification detected in a *Plasmodium falciparum* outbreak"

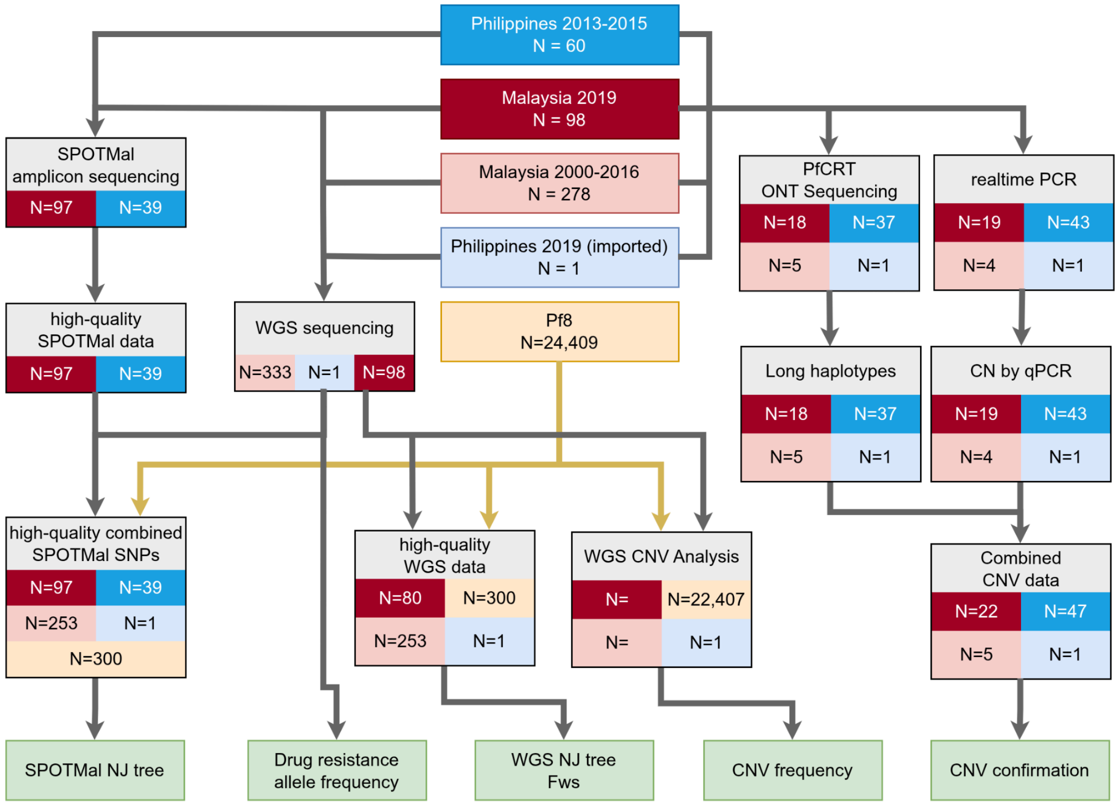


**Supplementary Figure 1. Flow chart summarising sample provenance for the analyses conducted in the study.** Background color indicates the country of the samples.

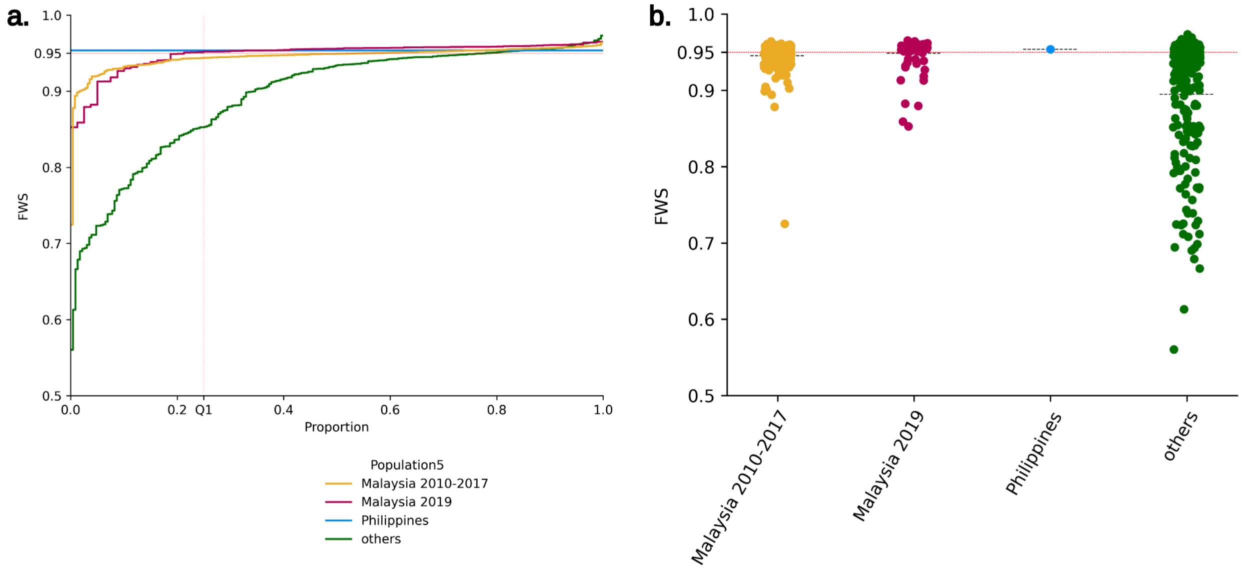


**Supplementary Figure 2. Within-sample diversity profiles in Malaysia and comparator populations.** The Fws was calculated using moimix with 116,704 biallelic SNPs (Malaysia 2010-2017 N=253, Malaysia 2019 N=80, Philippines N=1, other regions N=300). Panel a) presents the Fws distribution as Empirical Cumulative Distribution Function (eCDF) plot, and panel b) presents the distribution in categorical scatter plots . The complexity of Malaysian 2019 samples was comparable to Malaysia 2010-2017, indicating that transmission dynamics in Malaysia were relatively similar during that time period.

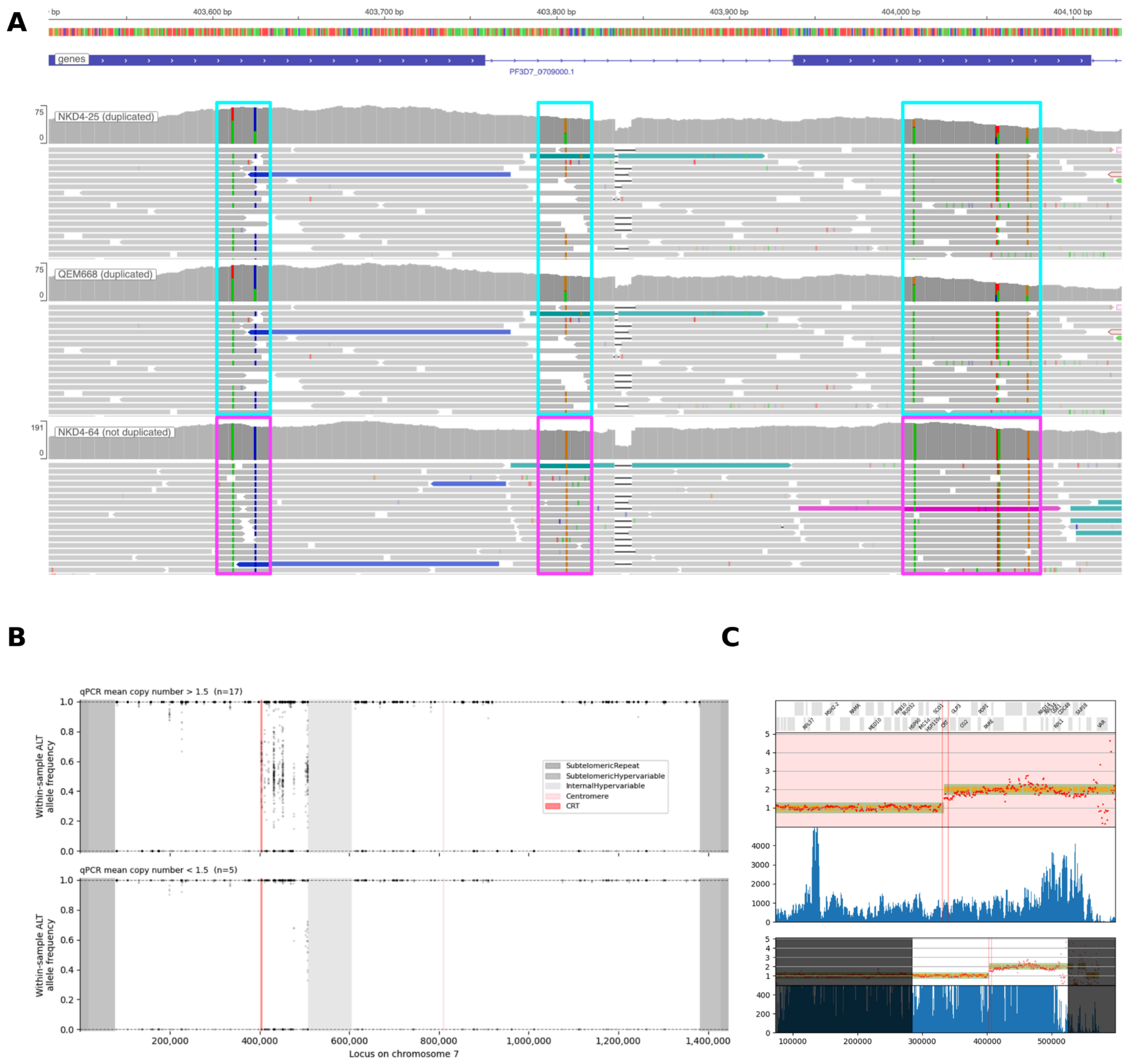


**Supplementary Figure 3. WGS evidence of the duplicated region encompassing *pfcrt***. Panel A shows a plot generated with IGV software illustrating two clonal samples with duplications (top and middle) and one sample with no duplication (bottom). Heterozygous SNPs within *pfcrt* are highlighted in cyan, while homozygous SNPs are highlighted in magenta. Panel B (upper) shows the within-sample allele frequency (WSAF) of alternate (ALT) alleles at all heterozygous loci on chromosome 7, across 17 independent, monoclonal ($F_{WS}$>0.95) Banggi Island (Malaysia 2019) samples with qPCR-based evidence of *pfcrt* duplication. A number of SNPs with WSAF around 0.5 are observed within f3D7_07_v3:403,612-507,272 demarking the approximate break points of the duplicated region. Panel B (lower) presents WSAF of ALT alleles in 5 independent, monoclonal samples with qPCR-based data indicative of single copies: the heterozygote signature observed in a) is absent among these samples. Panel C shows an example of a diagnostic plot generated for sample KF003 from running the MalariaGEN Pf8 CNV calling pipeline – the copy number state changes from 1 to 2 within the two red vertical lines which represent the boundaries of the *pfcrt* gene.
