## Supplementary tables for "*Pfcrt* copy number amplification detected in a *Plasmodium falciparum* outbreak"

**Supplementary Table 1. Within-sample diversity in Malaysia**

| FWS | Malaysia 2019 | Malaysia 2010-2017 |
|---|---|---|
| N | 80 | 253 |
| Mean | 0.949 | 0.945 |
| Median | 0.956 | 0.949 |
| Q1 | 0.952 | 0.944 |
| Q3 | 0.958 | 0.953 |
| N (FWS > 0.95) | 63 | 114 |
| N (FWS > 0.90) | 76 | 249 |

Data was generated on 333 independent Malaysian *P. falciparum* samples using 116,704 biallelic SNPs.

**Supplementary Table 2. Summary of genotype calls at drug resistance-associated variants in samples collected from the Philippines in 2013-2018.**

| Gene | Variant | Prop. Alt. Allele | No. Samples | No. Het. | Drug Assoc. | WHO Status |
|---|---|---|---|---|---|---|
| ***Pfcrt*** | 47V>47M | 2.56% | 39 | 1 | | |
| | 72C>72S | 7.69% | 39 | 3 | CQ | Validated |
| | 76K>76T | 7.69% | 39 | 3 | CQ | Validated |
| | 78F>78L | 2.56% | 39 | 1 | | |
| | 220A>220S | 2.50% | 40 | 1 | | |
| | 273H>273N | 41.03% | 39 | 10 | | |
| | 280W>280R | 2.56% | 39 | 1 | | |
| | 319F>319L | 2.50% | 40 | 1 | | |
| | 326N>326D | 7.50% | 40 | 3 | | |
| | 333T>333A | 2.50% | 40 | 1 | | |
| | 356I>356L | 2.56% | 39 | 1 | PPQ | Candidate |
| | 371R>371G | 2.56% | 39 | 1 | | |
| | 372E>372K | 2.56% | 39 | 1 | | |
| | 378F>378S | 2.56% | 39 | 1 | | |
| ***Pfmdr1*** | 86N>86Y | 2.50% | 40 | 0 | | |
| | 90D>90G | 2.50% | 40 | 1 | | |
| | 115C>115Y | 2.50% | 40 | 1 | | |
| | 117D>117N | 2.50% | 40 | 1 | | |
| | 130E>130K | 7.50% | 40 | 3 | | |
| | 977E>977K | 5.00% | 40 | 2 | | |
| | 1022R>1022G | 2.50% | 40 | 1 | | |
| | 1025I>1025N | 2.50% | 40 | 1 | | |
| | 1245R>1245K | 5.41% | 37 | 2 | | |
| | 1261F>1261S | 2.70% | 37 | 1 | | |
| | 1269I>1269V | 2.78% | 36 | 1 | | |
| | 1277T>1277I | 2.78% | 36 | 1 | | |
| ***Pfk13*** | 683I>683V | 2.56% | 39 | 1 | | |

| | | | | | | |
|---|---|---|---|---|---|---|
| | 678L>678M | 2.56% | 39 | 1 | | |
| | 676A>676T | 2.56% | 39 | 1 | | |
| | 671M>671I | 2.56% | 39 | 1 | | |
| | 663L>663Q | 2.56% | 39 | 1 | | |
| | 638G>638A | 2.56% | 39 | 1 | | |
| | 567E>567G | 2.50% | 40 | 1 | | |
| | 566V>566L | 2.50% | 40 | 1 | | |
| | 515R>515G | 7.50% | 40 | 3 | | |
| | 454V>454L | 2.50% | 40 | 1 | | |
| ***Pfdhfr*** | 15C>15R | 2.56% | 39 | 1 | | |
| | 20V>20I | 2.56% | 39 | 1 | | |
| | 21E>21K | 2.56% | 39 | 1 | | |
| | 25E>25K | 2.56% | 39 | 1 | | |
| | 29N>29S | 2.56% | 39 | 1 | | |
| | 54D>54N | 2.50% | 40 | 1 | | |
| | 59C>59R | 53.85% | 39 | 8 | PYR | Candidate |
| | 59C>59Y | 2.56% | 39 | 1 | | |
| | 68S>68L | 2.56% | 39 | 1 | | |
| | 108S>108N | 28.21% | 39 | 4 | PYR | Candidate |
| | 109W>109R | 5.13% | 39 | 2 | | |
| ***Pfdhps*** | 410E>410D | 2.56% | 39 | 1 | | |
| | 421M>421V | 2.56% | 39 | 1 | | |
| | 423N>423S | 2.56% | 39 | 1 | | |
| | 424E>424K | 2.56% | 39 | 1 | | |
| | 426A>426V | 2.56% | 39 | 1 | | |
| | 437G>437A | 76.92% | 39 | 3 | SP | Candidate |
| | 461K>461E | 2.56% | 39 | 1 | | |
| | 511P>511L | 2.63% | 38 | 1 | | |

Samples with a minimum read depth ≥5 at variant position; Number of samples with a heterozygous call at the variant position (No Het). *Prop. Alt. Allele = Proportion of samples carrying an alternate allele with a read depth ≥5; No. Het. = Number of samples with heterozygous alleles at that position.

### Supplementary Table 3. Combined results of realtime qPCR assay and *pfcrt* ONT long sequencing for Malaysian and Philippines samples.

| Country | Sample | Realtime qPCR assay | | | PfCRT ONT Haplotype Sequencing | | | CN | Remark |
|---|---|---|---|---|---|---|---|---|---|
| | | Avg FC | SD FC (range) | CN | Total depth | Estimated ratio over total depth | Haplotype | | |
| | KF1 | 0.9844 | 0.0075 (0.9776-0.9912) | 1 | 7281 | 0.827 | H2 | 1 | |
| | NKD4-101 | 0.9085 | 0.0181 (0.8920-0.9250) | 1 | 973 | 0.793 | H2 | 1 | |
| | NKD4-111 | - | - | - | 35 | 0.343 | H2 | 1 | Low quality sample |
| | NKD4-115 | 0.9083 | 0.0147 (0.8949-0.9217) | 1 | 4330 | 0.78 | H2 | 1 | |
| | NKD4-156 | 1.9462 | 0.0344 (1.9148-1.9777) | 2 | | | | | |
| | NKD4-167 | 0.9863 | 0.0106 (0.9766-0.9960) | 1 | 5058 | 0.835 | H2 | 1 | |
| | NKD4-183 | 1.8358 | 0.0002 (1.8356-1.8360) | 2 | | | | | |
| | NKD4-23 | 1.9462 | 0.1130 (1.8430-2.0493) | 2 | | | | | |
| | NKD4-56 | 2.1851 | 0.0711 (2.1201-2.2500) | 2 | | | | | |
| | NKD4-64 | 1.0459 | 0.0021 (1.0440-1.0479) | 1 | 6213 | 0.761 | H2 | 1 | |
| | QEM969 | 0.995 | 0.0022 (0.9930-0.9970) | 1 | 8430 | 0.798 | H1a | 1 | |
| | KF3 | 2.096 | 0.0205 (2.0772-2.1147) | 2 | 7394 | 0.417 | H1b | 2 | |
| | | | | | | 0.295 | H2 | | |
| | NKD4-103 | 1.7494 | 0.0261 (1.7256-1.7732) | 2 | 8371 | 0.379 | H1b | 2 | |
| | | | | | | 0.373 | H2 | | |
| | NKD4-114 | - | - | - | 37 | 0.333 | H2 | 1 | Low quality sample |
| | NKD4-118 | - | - | - | 41 | 0.354 | H1b | 2 | |
| | | | | | | 0.463 | H2 | | Low quality sample |
| | NKD4-154 | 1.7908 | 0.0546 (1.7410-1.8407) | 2 | 528 | 0.4 | H1b | 2 | |
| | | | | | | 0.438 | H2 | | |
| | NKD4-179 | 1.8611 | 0.0150 (1.8475-1.8748) | 2 | 5995 | 0.415 | H1b | 2 | |
| | | | | | | 0.341 | H2 | | |
| | NKD4-22 | 2.0152 | 0.2415 (1.7948-2.2356) | 2 | 75 | 0.331 | H1b | 2 | |
| | | | | | | 0.513 | H2 | | Low quality sample |
| | NKD4-25 | 2.1313 | 0.0233 (2.1100-2.1526) | 2 | 262 | 0.431 | H1b | 2 | |
| | | | | | | 0.379 | H2 | | |
| | NKD4-37 | 1.7587 | 0.0336 (1.7280-1.7894) | 2 | 1759 | 0.374 | H1b | 2 | |
| | | | | | | 0.38 | H2 | | |
| | NKD4-43 | 1.9717 | 0.0310 (1.9434-2.0000) | 2 | 55 | 0.291 | H1b | 2 | |
| | | | | | | 0.455 | H2 | | Low quality sample |
| | NKD4-70 | 1.8722 | 0.0151 (1.8584-1.8860) | 2 | 4033 | 0.374 | H1b | 2 | |
| | | | | | | 0.365 | H2 | | |
| | NKD4-79 | 2.0411 | 0.1072 (1.9433-2.1390) | 2 | 785 | 0.499 | H1b | 2 | |
| | | | | | | 0.363 | H2 | | |
| | NKD4-83 | 1.8655 | 0.1167 (1.7590-1.9721) | 2 | 11385 | 0.378 | H1b | 2 | |
| | | | | | | 0.292 | H2 | | |
| | NQE4-28 | 1.9628 | 0.0315 (1.9340-1.9916) | 2 | 5441 | 0.42 | H1b | 2 | |
| | | | | | | 0.333 | H2 | | |
| | QEM615 | - | - | - | 138 | 0.349 | H1b | 2 | |
| | | | | | | 0.43 | H2 | | |
| | QEM668 | 1.8389 | 0.0656 (1.7790-1.8988) | 2 | 2594 | 0.377 | H1b | 2 | |
| Malaysia | | | | | | 0.407 | H2 | | |
| | PF-2013-03 | 0.955 | 0.0055 (0.9500-0.9600) | 1 | 1292 | 0.854 | H1b | 1 | |
| | Pf-2013-04 | 0.89 | 0.0219 (0.8700-0.9100) | 1 | | | | | |
| | Pf-2013-09 | 0.95 | 0.0329 (0.9200-0.9800) | 1 | | | | | |
| | Pf-2013-11 | 0.965 | 0.0712 (0.9000-1.0300) | 1 | | | | | |
| | PF-2013-17 | 0.905 | 0.0383 (0.8700-0.9400) | 1 | 813 | 0.924 | H1a | 1 | |
| Philippines | Pf-2013-28 | 0.92 | 0.0219 (0.9000-0.9400) | 1 | | | | | |

| | | | | | | | | | |
|---|---|---|---|---|---|---|---|---|---|
| | Pf-2013-30 | 1.12 | 0.2300 (0.9100-1.3300) | 1 | | | | | |
| | PF-2013-40 | 1.01 | 0.0438 (0.9700-1.0500) | 1 | 170 | 0.191 | H1a | 1 | Probably polyclonal; asymmetric haplotype ratio |
| | | | | | | 0.709 | H1b | | |
| | Pf-2013-57 | 0.895 | 0.2027 (0.7100-1.0800) | 1 | | | | | |
| | Pf-2013-61 | 0.985 | 0.0274 (0.9600-1.0100) | 1 | | | | | |
| | Pf-2013-73 | 1.01 | 0.0329 (0.9800-1.0400) | 1 | 17 | 0.882 | H1a | 1 | Low quality sample |
| | PF-2013-75 | 1.025 | 0.1150 (0.9200-1.1300) | 1 | 125 | 0.904 | H1a | 1 | |
| | PF-2013-78 | 0.925 | 0.0164 (0.9100-0.9400) | 1 | 17697 | 0.869 | H1a | 1 | |
| | PF-2013-83 | 1.05 | 0.0548 (1.0000-1.1000) | 1 | 974 | 0.923 | H1a | 1 | |
| | PF-2015-11 | 0.86 | 0.0000 (0.8600-0.8600) | 1 | 281 | 0.826 | H1b | 1 | |
| | PF-2015-12 | 0.92 | 0.0548 (0.8700-0.9700) | 1 | 742 | 0.906 | H1a | 1 | |
| | PF-2015-15 | 1.195 | 0.1917 (1.0200-1.3700) | 1 | 56 | 0.875 | H1a | 1 | Low quality sample |
| | PF-2015-16 | 0.995 | 0.0493 (0.9500-1.0400) | 1 | 1413 | 0.89 | H1b | 1 | |
| | PF-2015-18 | 0.86 | 0.0000 (0.8600-0.8600) | 1 | 697 | 0.898 | H1a | 1 | |
| | PF-2015-19 | 0.935 | 0.0822 (0.8600-1.0100) | 1 | 178 | 0.362 | H1a | 1 | single-copy pfcrt with minor sequence variation |
| | | | | | | 0.537 | H1b | | |
| | PF-2015-20 | 0.985 | 0.0383 (0.9500-1.0200) | 1 | 1652 | 0.883 | H1a | 1 | |
| | PF-2015-21 | 1.035 | 0.0164 (1.0200-1.0500) | 1 | 40 | 0.9 | H1a | 1 | |
| | PF-2015-22 | 0.935 | 0.0712 (0.8700-1.0000) | 1 | 462 | 0.92 | H1a | 1 | |
| | PF-2015-39 | 0.72 | 0.0767 (0.6500-0.7900) | 1 | 15 | 0.933 | H1a | 1 | Low quality sample |
| | PF-2015-42 | 1.08 | 0.1315 (0.9600-1.2000) | 1 | 17 | 0.882 | H1a | 1 | Low quality sample |
| | PF-2015-43 | 1.105 | 0.2136 (0.9100-1.3000) | 1 | 5931 | 0.897 | H1b | 1 | |
| | PF-2015-44 | 1.015 | 0.0931 (0.9300-1.1000) | 1 | 227 | 0.899 | H1a | 1 | |
| | PF-2015-46 | 0.965 | 0.0602 (0.9100-1.0200) | 1 | 2542 | 0.852 | H1a | 1 | |
| | PF-2015-48 | 0.995 | 0.0164 (0.9800-1.0100) | 1 | 40 | 0.9 | H1a | 1 | Low quality sample |
| | PF-2015-49 | 1.085 | 0.1260 (0.9700-1.2000) | 1 | 113 | 0.823 | H1b | 1 | |
| | PF-2015-50 | - | - | - | 24 | 0.917 | H1a | 1 | Low quality sample |
| | Pf-2015-58 | 1.095 | 0.1807 (0.9300-1.2600) | 1 | | | | | |
| | PF-2015-59 | 1.05 | 0.1095 (0.9500-1.1500) | 1 | 229 | 0.572 | H1b | 1 | |
| | PF-2015-76 | 0.98 | 0.1643 (0.8300-1.1300) | 1 | 50 | 0.94 | H1a | 1 | |
| | PF-2015-80 | 1.105 | 0.1917 (0.9300-1.2800) | 1 | 112 | 0.821 | H1b | 1 | |
| | PF-2015-81 | 0.895 | 0.0055 (0.8900-0.9000) | 1 | | | | | Low quality sample |
| | PF-2015-82 | 1.13 | 0.1862 (0.9600-1.3000) | 1 | 1644 | 0.918 | H1a | 1 | |
| | PF-2017-18-36 | - | - | - | 942 | 0.914 | H1a | 1 | |
| | PF-2018-15 | 1.04 | 0.0657 (0.9800-1.1000) | 1 | 2697 | 0.916 | H1a | 1 | |
| | Pf-2018-28 | 1.135 | 0.2574 (0.9000-1.3700) | 1 | | | | | |
| | PF-2018-37 | | | | 2382 | 0.903 | H1a | 1 | |
| | PF-2018-52 | | | | 9623 | 0.901 | H1a | 1 | |
| | PF-2018-60 | 0.965 | 0.0164 (0.9500-0.9800) | 1 | 62 | 0.903 | H1a | 1 | Low quality sample |
| | PF-2018-64 | 0.93 | 0.0329 (0.9000-0.9600) | 1 | 3729 | 0.894 | H1a | 1 | |
| | Pf-2018-8 | 0.92 | 0.0219 (0.9000-0.9400) | 1 | | | | | |
| | PF-2013-72 | 2.11 | 0.0329 (2.0800-2.1400) | 2 | 674 | 0.591 | H1b | 2 | |
| | | | | | | 0.305 | H3 | | |
| | PF-2015-47 | 1.95 | 0.1643 (1.8000-2.1000) | 2 | 165 | 0.255 | H1a | 2 | |
| | | | | | | 0.639 | H2 | | |
| | QEM192* | 2.0389 | 0.1084 (1.9400-2.1379) | 2 | 5112 | 0.32 | H1b | 2 | |
| | | | | | | 0.287 | H4 | | |

*Imported case to Malaysia (2011), with travel history to the Philippines.

**Supplementary Table 4. Summary of the demography of Malaysian 2019 outbreak cases.**

| Site | Country | Collection period | Age category (years) | | | % Male patients |
|---|---|---|---|---|---|---|
| | | | <5 | 5-15 | >15 | |
| Sabah | Malaysia | Jan 2019 - June 2023 | 11/109 (10.09%) | 61/109 (55.96%) | 37/1409 (33.94%) | 62/109 (56.88%) |

**Supplementary Table 5. TaqMan *pfcrt* copy number quantitation assay primer and probe sequences and *pfcrt* ONT sequencing primers**

| Primer label | Sequences (5’-3’) |
|---|---|
| Pfcrt_TaqMan_F | ACGACACCGAAGCTTTAATTTAC |
| Pfcrt_TaqMan_R | TTTCCAGTAGTTCTTGTAAGACCT |
| Pfcrt_TaqMan_Probe | TGCTATATCCATGTTAGATGCCTGTTCAGT |
| B-tubulin_Taqman_F | CCCATTCCCACGTTTACATTTC |
| B-tubulin_Taqman_R | GGCACAGTTAAGGCTCTGTAT |
| B-tubulin_Taqman_Probe | CGGGTTTGCTCCTTTAACTAGTAGAGGC |
| Pfcrt_ONT_F | CCGTTAATAATAAATACACGCAG |
| Pfcrt_ONT_R | TCCTTATAAAGTGTAATGCGATAGC |
